# Spurious inference when comparing networks

**DOI:** 10.1101/619957

**Authors:** Damien R. Farine, Lucy M. Aplin

## Abstract

Comparing networks is challenging. Social networks can be shaped by many factors. Failing to adequately consider non-social processes, including sampling artefacts, can lead to spurious conclusions about differences in social networks among groups. Here we demonstrate that incorrect application of statistical testing methods when comparing networks can generate very high rates of false positives. We then show that null models, specifically pre-network permutation tests, can control for non-social differences in networks and substantially reduce rates of false positives.

## Main text

Comparing networks is challenging (1–3). van Leeuwen *et al.* (4) (*herein* vLCH) use variation in networks among four chimpanzee groups as evidence for cultural differences in social behaviour. However, social networks can be shaped by many factors. The authors control for group size, an obvious correlate of many network metrics, but do not adequately consider alternative processes that can generate differences in social networks among groups. Many are unrelated to social decisions. For example, habitat geometry can maintain highly-repeatable between-generation network community structure in wild populations of songbirds (5).

We illustrate why vLCH’s study cannot be used as evidence for culturally-driven differences among chimpanzee groups using simulations (see 6). Briefly, we randomly placed N group members (with group sizes matching vLCH’s data) in an arena by drawing X and Y coordinates from a uniform distribution (forming an area matching each group’s enclosure). We then calculated inter-individual distances, using 1 unit (relative to side length) as our measure of ‘being connected’ and a chain rule to define parties (repeating this process 10 times to generate 10 observations per group). We applied the statistical testing approaches described by vLCH—constructing models with and without group identity as random effects, and using post-network permutation tests. We repeated this procedure 100 times to estimate Type I error rates. Our aim was not to replicate exactly the data produced by vLCH’s observations, but instead to show that simple non-social processes can generate highly significant and spurious results when using incorrect hypothesis-testing methods. Our results confirm this; every test performed by vLCH found significant differences between groups, with their methods generating anti-conservative P values and extremely high rates of false positives (~94.3%, Table 1a).

**Table 1.** Summary of the simulation results. Lower P and Upper P represent the 2.5% and 97.5% quantiles of the P value distribution respectively. % False Positives is the proportion of simulations that generated P values that were significant at different significance thresholds.

| Test | (a) van Leeuwen et al. method |  |  |  | (b) Pre-network permutation method |  |  |  |
| --- | --- | --- | --- | --- | --- | --- | --- | --- |
|  | Lower P | Upper P | % False Positives (P < 0.05) | % False Positives (P < 0.001) | Lower P | Upper P | % False Positives (P < 0.05) | % False Positives (P < 0.001) |
| Party size | 4.37x10<sup>-13</sup> | 0.017 | 98 | 91 | 0.232 | 0.99 | 0 | 0 |
| Proximity | 2.44x10<sup>-29</sup> | 1.58x10<sup>-6</sup> | 100 | 100 | 0.0143 | 0.995 | 6 | 3 |
| Strength | 0 | 0.001 | 100 | 99 | 0 | 0.990 | 11 | 4 |
| Eigen. Cent. | 0 | 0.009 | 100 | 52 | 0.015 | 0.990 | 7 | 2 |
| Reach | 0 | 0 | 100 | 100 | 0.005 | 1 | 9 | 3 |
| Clustering | 0 | 0.03 | 98 | 83 | 0.049 | 0.985 | 3 | 2 |
| Affinity | 0 | 0 | 100 | 100 | 0.020 | 1 | 10 | 1 |

While comparison across networks remains challenging, one solution is to use permutation tests (3). Specifically, pre-network permutation tests that swap observations among individuals within groups can be used generate realistic ‘randomised’ networks, and thus produce a distribution of expected test statistics in which the social process of interest has been removed (7–9). Post-network (or node) permutations are not useful for comparing networks because they do not change the network-level distribution of metrics, making them only useful for within-network contrasts (8). Applying pre-network permutation tests on exactly the same simulated data produced reliable Type I errors rates (~3.6%, Table 1b), because pre-network permutations effectively controlled for a simple non-social causal effect in our simulated data: density. We note, however, that density is unlikely to be the sole driver of between-group difference found by vLCH.

We conclude that as it stands, the study by vLCH does not provide robust evidence for significant cultural differences in social behaviour. Using appropriate hypothesis-testing methods completely changes the conclusions of vLCH’s study. More broadly, even if significant between-group differences remain, attributing them to culture requires following groups over longer time scales in order to rule out alternative mechanisms arising from interactions between particular individuals. Individual differences in personality are well-known to shape group-level collective behaviours (10), but this doesn’t necessarily lead to cultures that outlast the individuals involved.

## Acknowledgements

DRF and LMA were funded by the Max Planck Society and by the DFG Centre of Excellence 2117 “Centre for the Advanced Study of Collective Behaviour” (ID: 422037984). We thank two anonymous reviewers for constructive comments that have improved our letter.

